# Clonal analysis by tunable CRISPR-mediated excision

**DOI:** 10.1101/394221

**Authors:** Anna F. Gilles, Johannes B. Schinko, Magdalena I. Schacht, Camille Enjolras, Michalis Averof

## Abstract

Clonal marking techniques based on the Cre/lox and Flp/FRT systems are widely used in multicellular model organisms to mark individual cells and their progeny, in order to study their morphology, growth properties and developmental fates. The same tools can be adapted to introduce specific genetic changes in a subset of cells within the body, i.e. to perform mosaic genetic analysis. Marking and manipulating distinct cell clones requires control over the frequency of clone induction, which is sometimes difficult to achieve. Here we present Valcyrie, a new method that replaces the conventional Cre or Flp recombinase-mediated excision of a marker cassette by CRISPR-mediated excision. A major advantage of this approach is that CRISPR efficiency can be tuned in a predictable fashion by manipulating the degree of sequence complementarity between the CRISPR guide RNA and its targets. We establish the method in the beetle *Tribolium castaneum*. We demonstrate that clone marking frequency can be tuned to generate embryos carrying single marked clones. The Valcyrie approach can be applied to a wide range of experimental settings, for example to modulate clone frequency with existing tools in established model organisms and to introduce clonal analysis in emerging experimental models.

## Introduction

Clonal analysis originates from efforts to track the fate of individual cells and their progeny during development (reviewed in Kretzschmar and Watt, 2012). Early studies used cell transplantation from genetically distinct donors or labelling using vital dyes to track the fate of marked cells. Spontaneous somatic mutations producing a visible cellular phenotype, such as the loss of pigmentation, were also used to observe the territories occupied by cell clones in mosaic animals and plants. In principle, any heritable somatic change associated with a visible marker can be used for this purpose, and different types of genetic events have been exploited for this, including chromosome loss, transposon excision, genetic rearrangement and mitotic recombination (Garcia-Bellido et al., 1973; Janning, 1978; Nicolas et al., 1996; Stern, 1936; Vincent et al., 1995).

Clonal analysis received a great boost with the introduction of recombinases and fluorescent proteins in the 1990s: recombinases made the task of inducing genetic rearrangements more efficient and easier to engineer (Golic and Lindquist, 1989; Orban et al., 1992), and fluorescent proteins provided versatile markers for visualising the cell clones. The most common method for marking cell clones, used until today, consists of expressing a recombinase (bacteriophage-derived Cre or yeast-derived Flp) in the cells of interest, where it induces recombination among pairs of the respective target site (*lox* or *FRT*). These target pairs can be placed either in tandem, to excise the intervening DNA fragment containing a marker (flip-out), or on homologous chromosomes, to exchange chromosome arms carrying markers and/or mutations. Many elaborations of those two configurations have been used, varying in the number and complexity of genetic rearrangements (reviewed in Griffin et al., 2014; Richier and Salecker, 2014).

Today, clonal marking techniques are widely applied to visualise the morphology and behaviour of single cells within complex tissues, and to investigate lineage relationships, stem cell dynamics and tissue growth (e.g. Blanpain and Simons, 2013; Chen et al., 2016; Kuchen et al., 2012; Livet et al., 2007; Morante and Desplan, 2008; Snippert et al., 2010).

The same tools have also been adapted to create genetic mosaics where specific genetic changes are introduced in some cells of the body. Such genetic mosaics have been invaluable for discovering the cells where a gene function is needed, for distinguishing a gene’s cell autonomous and non-autonomous functions, and for studying the effects of mutations that would be lethal in the entire organism (e.g. Hotta and Benzer, 1972; Lee and Luo, 1999; Rajewsky et al., 1996). By allowing us to juxtapose cells of different genotypes in a tissue, mosaics have allowed us to discover the genes that underpin key cell-cell interactions during development and tissue homeostasis (e.g. Brook et al., 1996; Meyer et al., 2014).

Clonal analysis techniques require a good control over the frequency of clone induction. If the frequency is too low, a very large number of individuals must be screened in order to find suitable cell clones to analyse and thus experiments become very laborious. On the other hand, if the frequency is too high, independent cell clones frequently merge with each other and thus information on the clonal relationships of cells is lost. In established model organisms, such as *Drosophila* and mice, the research community has developed and fine-tuned these techniques over decades, generating tools with clone induction frequencies that are well suited for the purpose. When establishing these tools in new experimental models, however, adjusting these frequencies can be laborious and frustrating. Here we present the difficulties that we faced while establishing clonal analysis tools in the beetle *Tribolium castaneum*. We show that the Cre/lox system has such high activity in *Tribolium* that a large number of marked cell clones are generated even by very low (undetectable) levels of Cre expression, driven by the leaky activity of a heat-shock promoter.

To overcome this problem and to optimise the frequency of induction of marked clones, we developed an alternative method for generating marked clones, based on the excision of a marker cassette by the CRISPR gene editing system (Cong et al., 2013; Gilles and Averof, 2014; Jinek et al., 2012; Mali et al., 2013), rather than by Cre- or Flp-mediated recombination (Figure 1). This approach, which we name Valcyrie (for <u>V</u>ersatile and <u>a</u>djustable labelling of <u>c</u>lones b<u>y</u> random CRISPR-induced <u>e</u>xcision), offers the major advantage that the efficiency of clone induction can be adjusted by manipulating the degree of sequence complementarity between the CRISPR guide RNA (gRNA) and its targets. We demonstrate the effectiveness of this approach in *Tribolium*. In established model organisms, this tunable method can be readily applied on existing flip-out lines established with constructs carrying *lox, FRT* or *attP/B* recombinase sites.

**Figure 1.**
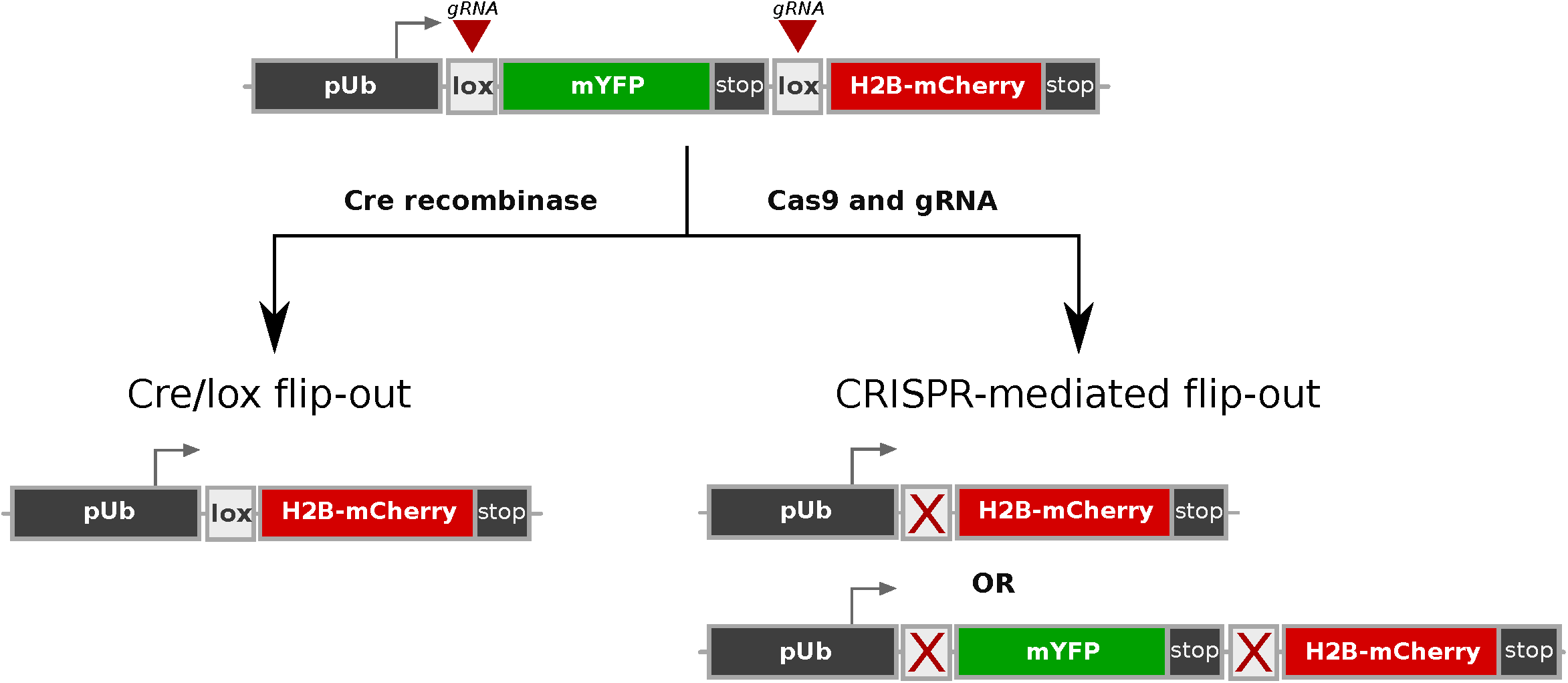
Recombinase-and CRISPR-mediated approaches for clonal analysis. In conventional recombinase-mediated approaches, the action of a recombinase drives a recombination event that stably marks the cell and its clonal progeny. Here, the Cre recombinase acts on a pair of *lox* sites, causing the excision of mYFP and bringing H2B-mCherry under the control of the *pUb* promoter. In the CRISPR-mediated approach, the same excision is caused by double-strand DNA breaks generated by the action of Cas9 and gRNAs targeting the *lox* sites. The CRISPR-mediated approach can also generate point mutations, which prevent further CRISPR activity on the *lox* sites. The CRISPR-mediated approach allows us to control the timing and frequency of excision by manipulating the expression of Cas9 and the degree of sequence complementarity between the CRISPR gRNAs and their targets. Mutated lox sites are indicated by a red cross.

## Results and Discussion

### Activity and leakiness of the Cre/lox system in *Tribolium*

To establish the Cre/lox system for clonal analysis in *Tribolium castaneum*, we generated stable transgenic lines bearing conventional Cre and *lox* flip-out constructs. The Cre-expressing line (*Tc-hsp68::Cre*) carries the coding sequence of the Cre recombinase driven by the heat-inducible promoter of the *Tc-hsp68* gene (Schinko et al., 2012). The flip-out construct, *pUb::lox-mYFP-lox-H2BmCherry*, carries a flip-out cassette consisting of the coding sequence of membrane-localised Yellow Fluorescent Protein (mYFP) flanked by *loxN* sites (Livet et al., 2007), followed by a fusion of coding sequences for the fluorescent protein mCherry (Shaner et al., 2004) and histone H2B of *Drosophila* (H2B-mCherry); the flip-out construct is transcribed by the ubiquitous *pUb* promoter (see Materials and Methods). In the absence of a Cre transgene, embryos carrying the flip-out construct show ubiquitous expression of mYFP in late blastoderm and early germ band stages (Figure 2A).

**Figure 2.**
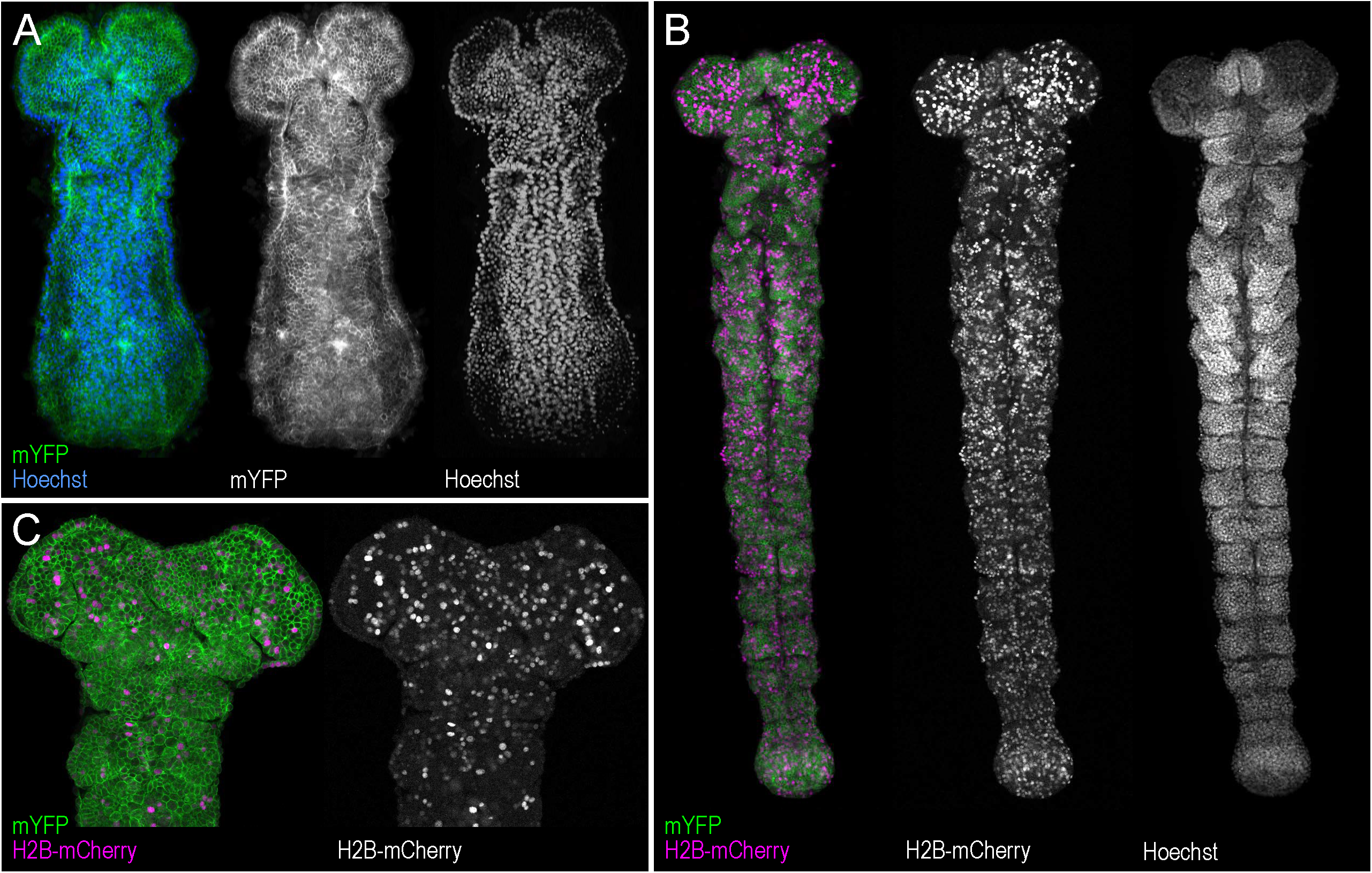
Leakiness of the Cre/lox system in *Tribolium*. **A.** Uniform expression of mYFP (in green) in *Tribolium* embryo carrying the *pUb::lox-mYFP-lox-H2BmCherry* flip-out construct, in the absence of a Cre transgene. The embryo was also stained with Hoechst 34580 to visualise all nuclei (in blue). The mYFP and Hoechst channels are shown separately at the right of the panel. **B-C.** mYFP and H2B-mCherry expression (green and magenta respectively) in *Tribolium* embryos carrying the same flip-out construct in the presence of the *Tc-hsp68::Cre* transgene. H2B-mCherry-marked nuclei are visible even in the absence of a heat shock to induce the expression of Cre. The H2B-mCherry channel and Hoechst 34580 staining (in panel B) are shown separately at the right of each panel. Panels show the entire germband (B) and head region (C) of separate embryos.

By crossing these transgenic lines, we obtained embryos carrying both the heat-inducible Cre and the flip-out construct. In these embryos, a large number of cells lacked mYFP expression and instead expressed H2B-mCherry, indicating that the *lox-mYFP-lox* cassette was efficiently excised in the presence of Cre (Figure 2B,C). We expected the excision to depend on the induction of Cre by a heat shock, but in fact we observed that embryos carry a large number of H2B-mCherry-expressing cells both with and without heat shock. These results suggest that sufficient Cre activity is present even in the absence of a heat shock, due to leaky activity of the *Tc-hsp68* promoter. In the absence of heat shock, Cre mRNA was not detectable by *in situ* hybridization, suggesting that very low levels of Cre are sufficient to mobilise the *lox-mYFP-lox* cassette. This Cre activity was present when Cre was provided either maternally or zygotically. Leaky activity of Cre has also been reported in similar experiments carried out in *Drosophila* (Frickenhaus et al., 2015).

### Design of Valcyrie

In standard Cre/lox flip-out constructs, the expression of Cre drives recombination of paired *lox* sites, excising a marker gene and bringing a sequence encoding a second marker or gene of interest under the control of a ubiquitous promoter. In the Valcyrie approach, the same flip-out construct can be used, but excision is mediated by a pair of double-strand breaks targeted to the *lox* sites by CRISPR (Figure 1). The induction of double-strand breaks is dependent on the expression of the Cas9 protein and a specific gRNA, which together constitute the CRISPR endonuclease (Hwang et al., 2013; Jinek et al., 2012). Once the CRISPR endonuclease has cleaved the two *lox* sites, the cell’s DNA repair mechanisms re-join the broken DNA ends; the cleaved sites may be repaired precisely, carry small mutations, or lack the entire fragment between the two targeted sites.

This design provides the opportunity to modulate the production of marked clones in different ways. First, the promoters used to express the Cas9 and the gRNA can influence when, where and at what frequency the marked clones are induced. Here, to test this approach, we used the *Tc-hsp68* promoter to drive strong expression of Cas9 upon heat shock, and two constitutive RNA polymerase III promoters (*Tribolium U6a* or *U6b*; Gilles et al., 2015) to drive the expression of gRNAs.

Second, the level of CRISPR activity can be tuned down by manipulating the degree of sequence complementarity between the gRNA and its targets on the flip-out construct. The CRISPR system is known to tolerate mismatches especially towards the 5’end of the gRNA (Hsu et al., 2013; Jiang et al., 2013; Fu et al., 2013). With this in mind, we designed and tested two sets of gRNAs expected to target the *lox* sites of our flip-out construct with different efficiencies. First, we designed a pair of gRNAs targeting each *lox* site as efficiently as possible (given the sequence constraints of the flip-out construct, see Materials and Methods): *gRNA Lox.R* fully matches one of the *lox* sites, and *gRNA Lox.L* matches the second *lox* site with a single nucleotide mismatch at the 5’ end of the gRNA (Figure 3B). Second, we designed a single gRNA that is expected to target both *lox* sites with a lower efficiency: *gRNA Lox.Uni* carries two mismatches with each target, located within the first 6 nucleotides of the gRNA (Figure 3B).

**Figure 3.**
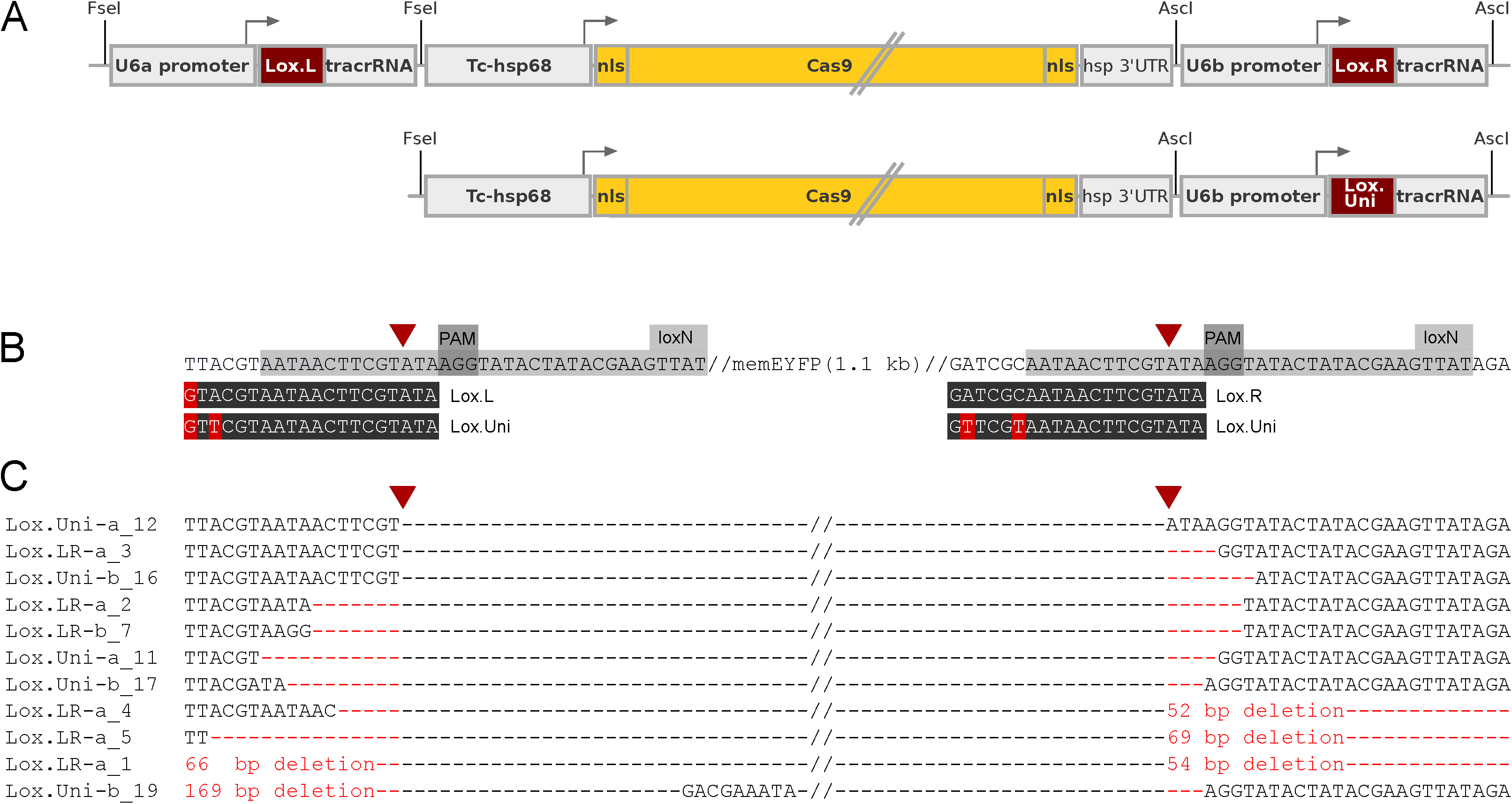
CRISPR-mediated targeting of *lox* sites. **A.** Schematic representation of the *Valcyrie.LR* and *Valcyrie.Uni* constructs (top and bottom, respectively). **B.** Sequence of the *loxN* sites present in the flip-out construct *pUb::lox-mYFP-lox-H2BmCherry* and the targeting sequences of the *Lox.L, Lox.R* and *Lox.Uni* gRNAs. CRISPR targeting requires the presence of a PAM motif (NGG) in the target sequence. The single PAM found in *loxN* sites constrains the design of the gRNAs, which must be placed at the edge on the *lox* site, encompassing 14 nucleotides of the *lox* site and 6 nucleotides of the sequence flanking each *lox* site. The mismatches between the target sites and the gRNAs are marked in red. The CRISPR cleavage site is indicated by an arrowhead. **C.** Sequences resulting from independent excision events (aligned to the target sequences). The excision events are usually imprecise, as expected for repair by non-homologous end joining (NHEJ). In four cases the deletions extended upstream or downstream of the targeted *lox* sites.

Besides generating pairs of double-strand breaks, leading to the excision of the *lox-mYFP-lox* cassette, we expect that in some cells CRISPR activity will generate point mutations (indels) in individual *lox* sites. The majority of these mutations will produce modified *lox* sequences that can no longer be targeted by CRISPR (Figure 1). As these mutations accumulate, they should lead to a decline and eventually an arrest of clone-marking excisions, producing mosaic animals where only a small number of cells have been marked, and the majority of cells remain unlabelled (carrying mutated *lox* sites). This would be beneficial for clonal analysis.

To facilitate the genetic crosses bringing the Cas9, gRNA and flip-out elements together, we cloned the Cas9 transgene and each set of gRNAs in the same vector (see Figure 3A) and generated two independent transgenic lines for each construct, termed *Valcyrie.LR #22* and *#39* (expressing the *Lox.L* and *Lox.R* gRNAs) and *Valcyrie.Uni #6* and *#11* (expressing the *Lox.Uni* gRNA). The lines carry transgene insertions that segregate as single Mendelian loci. *Valcyrie.LR #22, Valcyrie.Uni #6* and *Valcyrie.Uni #11* are kept as viable homozygous lines. The insertion present in *Valcyrie.LR #39* is homozygous lethal and it is therefore kept in heterozygous condition.

### Generating marked clones at adjustable frequencies via CRISPR-mediated excision

To test whether CRISPR-mediated excision is effective in generating marked clones, we crossed beetles from each of the *Valcyrie.LR* and *Valcyrie.Uni* lines to beetles carrying the flip-out construct *pUb::lox-mYFP-lox-H2BmCherry* (the same flip-out line that we previously tested for Cre-mediated excision). We collected mixed embryonic stages, from early to late germband stages and subjected them to a heat shock (46°C for 10 min, see Schinko et al., 2012) to induce the expression of Cas9. We let these embryos develop for an additional 12 hours at 32°C before fixation, and screened them for H2B-mCherry expression. As a control, we screened embryos from identical crosses that were not heat shocked, but were collected and fixed as described above. 10 to 25 embryos from each cross were screened by confocal microscopy.

We detected H2B-mCherry-positive cells in embryos from all four crosses with variable frequency after heat shock (Figure 4, Table 1). A higher proportion of embryos with visible clones was recorded with the *Valcyrie.LR #22* and *#39* lines (>80% of embryos carrying the construct) than with the *Valcyrie.Uni #6* and *#11* lines (20-70% of embryos carrying the construct). This result is in line with our expectation that nucleotide mismatches of increasing severity in the gRNAs will generate flip-out clones at lower frequencies. We did not find any H2B-mCherry-positive cells in the controls for the *Valcyrie.LR #22, Valcyrie.Uni #6* and *Valcyrie.Uni #11* crosses, but we detected some marked cells in the control for *Valcyrie.LR #39*, indicating leaky expression of Cas9 in this line, in the absence of a heat shock. These results indicate that most Valcyrie lines give us control over the induction of marked clones.

**Table 1.**
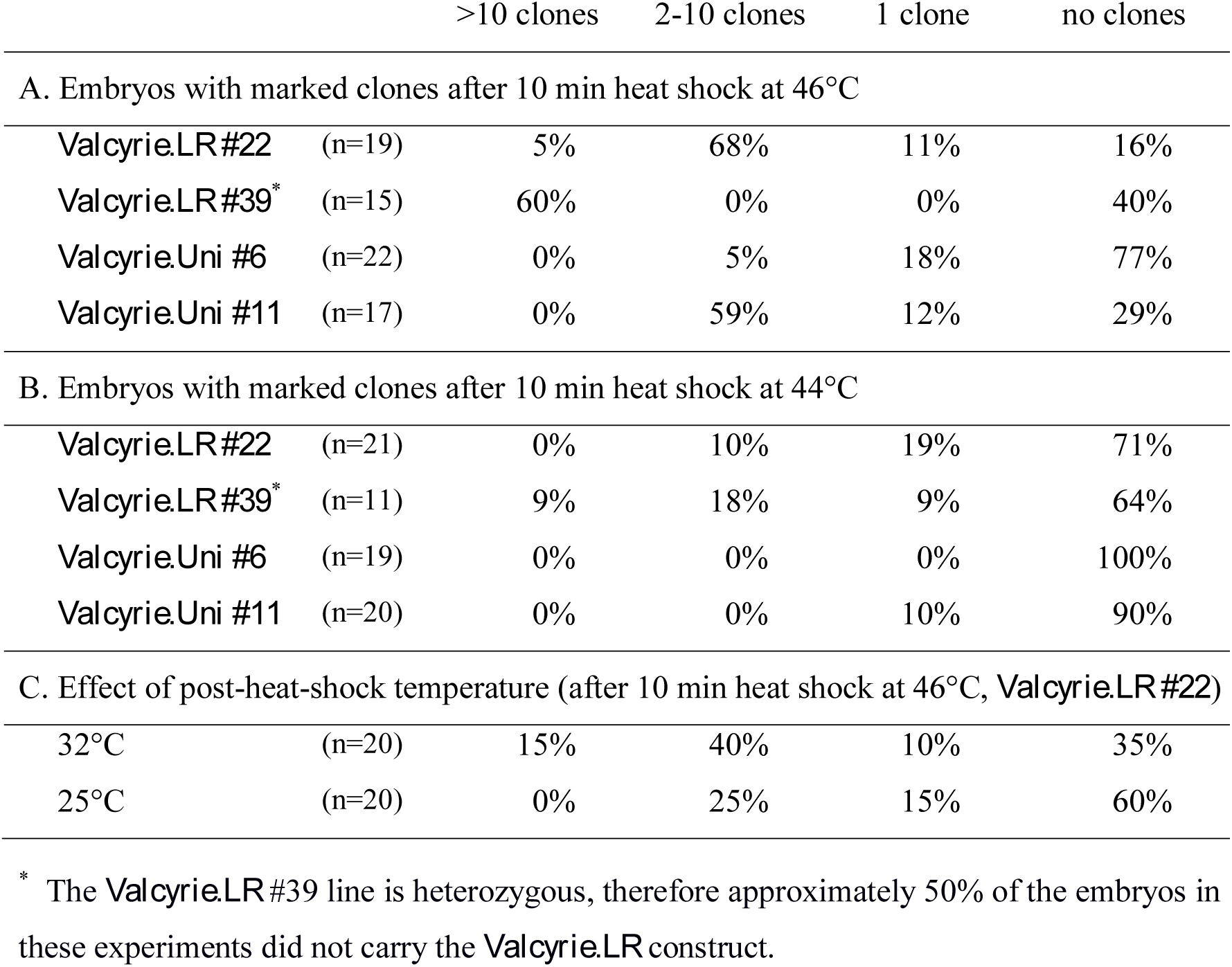
Frequency of marked cell clones. The number of distinct clusters of H2B-mCherry-expressing cells was scored per embryo. We classified embryos in four categories: embryos with more than 10 clusters per embryo, 2-10 clusters, one cluster, or none. Each cluster was taken to represent a cell clone.

**Figure 4.**
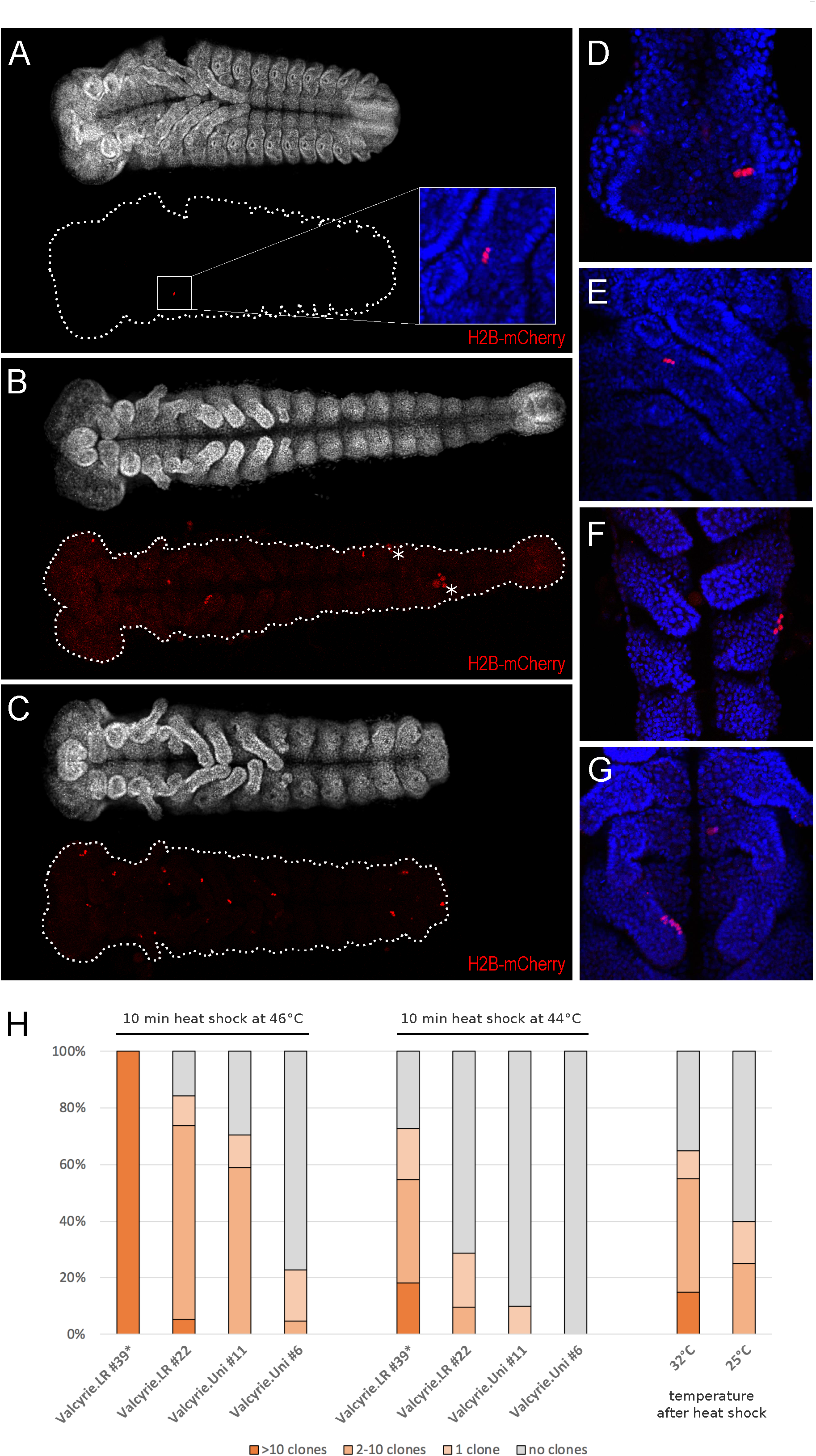
Cell clones marked by CRISPR-mediated excision in *Tribolium*. **A-C.** *Tribolium* embryos bearing cell clones marked by CRISPR-mediated excision, using Valcyrie. The top of each panel shows the morphology of the embryo by Hoechst 34580 staining, marking DNA in all nuclei. The bottom of each panel shows H2B-mCherry fluorescence (in red) marking the cell clones. Embryos were categorized as having one cell cluster (A), 2-10 cell clusters (B) or more than 10 clusters (C). Asterisks indicate autofluorescence in yolk cells. **D-G.** High magnification views of H2B-mCherry marked cell clones (in red) in different parts of the embryo, including posterior mesoderm (D), leg mesoderm (E), abdominal ectoderm (F), and leg ectoderm (G). Hoechst 34580 marks DNA in all nuclei (in blue). **H.** The frequency of marked clones is influenced by the sequence of the CRISPR gRNAs, the insertion site of the transgenes, the strength of the heat shock and the incubation temperature after the heat shock. *Valcyrie.LR* lines yield a higher number of clones per embryo than the *Valcyrie.Uni* lines, consistent with the degree of complementarity between the gRNAs and the *lox* targets (see Figure 3B). The frequency of clones can also be modulated by the heat shock inducing the expression of Cas9 (10 min at 44°C versus 46°C), or by the incubation temperature after the heat shock (25°C versus 32°C). The graphs are based on data shown in Table 1. The frequencies of the *Valcyrie.LR #39* line have been corrected considering that half of the embryos do not carry the Valcyrie transgenes.

In addition to scoring the presence or absence of H2B-mCherry-expressing cells, we also counted how many distinct clusters of H2B-mCherry-positive cells could be found in each embryo (Figure 4H, Table 1). We used the number of these clusters as a proxy of the number of independent cell clones marked in each embryo. As expected, embryos from the *Valcyrie.LR* lines have more clusters than embryos from the *Valcyrie.Uni* lines (Figure 4H), which we attribute to the different efficiency of the two sets of gRNAs. However, there is also a considerable difference between the two lines carrying each construct, which is likely due to different levels of Cas9 and gRNAs produced from each transgene insertion site.

The results presented so far show that introducing mismatches between the gRNAs and the *lox* target sites is an effective strategy for reducing the frequency of marked clones in a predictable fashion.

Based on previous work (Schinko et al., 2012), we expected that the temperature and/or the duration of the heat shock could influence the expression level of Cas9 and thus provide an additional means for tuning the frequency of clone labelling to desired levels. To investigate this possibility, we repeated the experiment described above with a milder heat shock, at 44°C instead of 46°C. As anticipated, we found that the number of marked cell clusters was greatly reduced with all four Valcyrie lines (Figure 4H, Table 1B).

Further, we suspected that the number of marked clones might also depend on the incubation temperature after the heat shock. To probe this, we collected embryos from *Valcyrie.LR #22* line at 32°C and subjected them to a heat shock for 10 minutes at 46°C. Then we let one batch of these embryos develop at 32°C for 20 hours, and another at 25°C for 40 hours. *Tribolium* development is approximately twice as fast at 32°C compared to 25°C, so these batches developed to comparable developmental stages. We found that the embryos kept at 32°C had more numerous marked cell clusters than the embryos kept at 25°C (Figure 4H, Table 1C). We speculate that, after the heat shock, CRISPR activity persists for longer at 32°C than 25°C.

### Distribution, timing, and nature of excisions

In these experiments we found that we were able to generate marked clones in all regions of the *Tribolium* embryo, including the anterior head, gnathal, thoracic and abdominal segments. Clones were visible in the ectoderm and the mesoderm of the germ band, in developing appendages and in the hindgut (Figure 4D-G), demonstrating that Valcyrie can be used to generate marked clones in most, if not all, cell populations of the *Tribolium* embryo.

We were particularly interested to determine whether we could induce flip-outs as early as in late blastoderm or early germband stages, because these are stages when important patterning events, such as establishing segmental boundaries and organ primordia, take place in *Tribolium* embryos. To answer that question, we subjected late blastoderm embryos (6-7 hours after egg laying, at 32°C) to a heat shock, and determined the onset of H2B-mCherry expression. We could detect the earliest H2B-mCherry-positive nuclei in early germ band stages (after 13 hours at 25°C), showing that Valcyrie constructs can be used to mark clones during these stages. However, we did not detect any H2B-mCherry expression in embryos that were subjected to a heat shock during earlier stages (5-6 hours after egg laying at 32°C, or younger). The timing of clone induction should correspond to the onset of heat-inducible expression (Schinko et al., 2012).

To molecularly characterize the excision events that take place in the flip-out construct, we extracted genomic DNA from a mix of embryos carrying flip-out clones, used PCR to amplify the region that lies between the paired *lox* sites, and sequenced several of these DNA fragments. We could detect deletions of variable length spanning that region (Figure 3C), as expected for sites repaired by non-homologous end joining. In two out of 11 sequenced clones, the deletions extended into the coding sequence of H2B-mCherry. In one other case, part of the *pUb* promoter was deleted. These results show that, while the majority of flip-outs generate marked clones, some Valcyrie excisions can also lead to the loss of both mYFP and H2B-mCherry marker expression.

### Conclusions

We have presented a novel method for generating marked cell clones, which relies on CRISPR. Compared with conventional recombinase-mediated clonal marking, our method provides additional means to control the frequency of clonal labelling, by manipulating CRISPR activity; this can be tuned in a predictable fashion by manipulating the degree of sequence complementarity between the CRISPR gRNAs and their targets. The method is highly versatile; the flexible design of gRNAs allows it to be applied on any target construct, including existing constructs with paired *lox, FRT* or *attP/B* sites, but also on new sites introduced through transgenes or on sites found naturally in the genome (see below).

We demonstrated the Valcyrie approach in the beetle *Tribolium castaneum*, an experimental model used for comparative studies of development and for studies on pest control. The genetic tools and resources available in this species have expanded significantly in recent years, to inducible gene expression, the GAL4/UAS system, a complete genome sequence, large RNAi and enhancer trapping screens, and CRISPR-mediated genome editing (Gilles et al., 2015; Richards et al., 2008; Schinko et al., 2012; Schinko et al., 2010; Schmitt-Engel et al., 2015; Trauner et al., 2009). However, clonal marking tools, which would allow us to mark and to track individual cell clones, have been lacking. Our initial effort to establish clonal analysis using a conventional Cre/lox flip-out approach was hampered by the leaky activity of the Cre recombinase, which led to uncontrolled clonal labelling. The Valcyrie approach allows us to overcome this problem, giving us control over the timing and frequency of clonal labelling.

Using Valcyrie, we have shown that the frequency of excision events can be tuned in several ways: by designing gRNAs having optimal or sub-optimal matches with their target sites (i.e. the *Valcyrie.LR* or the *Valcyrie.Uni* constructs), by choosing different transgene insertions, and by modulating the temperature of the heat shock and post-heat-shock treatments. Using the *Valcyrie.Uni #6* and *#11* lines, we have found conditions in which we reliably generated embryos that carry a single or a few marked clones. Additional levels of control can be envisaged by employing different promoters to drive the expression of Cas9 and/or gRNAs.

We applied Valcyrie to *lox* flip-out constructs, but the same approach could be used in any experiment that requires excision of a DNA fragment. In established model organisms it can be used on existing *FRT* or *lox* flip-out constructs, including constructs used for mosaic/clonal analysis of gene functions, to fine-tune the frequency of excision. In principle, CRISPR-mediated excision could also be extended to perform clonal analysis in the absence of flip-out transgenes, by targeting sequences already present in the genome. For example, target sites flanking a *cis*-regulatory element or an exon could be used to excise that element in specific cell populations (using an appropriate driver for Cas9) or in random cell clones. The major challenges in that case will be finding ways to mark the excision events and identifying CRISPR target sites that have no phenotype when individually mutated (e.g. targeting intergenic or intronic regions).

Our method of CRISPR-mediated clonal analysis may be of particular interest in experimental models where conventional Cre/lox- and Flp/FRT-based clonal analysis tools are not yet available, but where transgenesis and CRISPR have been established, such as *Nematostella vectensis* and *Parhyale hawaiensis* (Ikmi et al., 2014; Renfer and Technau, 2017; Pavlopoulos and Averof, 2005; Martin et al., 2016). The tunability of Valcyrie provides a straightforward way for adapting and optimising clonal analysis tools in diverse experimental settings and organisms.

## Materials and Methods

### Cre/lox and Valcyrie constructs

The *Tc-hsp68::Cre* construct was generated by cloning the Cre coding sequence under the heat-inducible promoter of the *Tc-hsp68* gene, including the endoHSE upstream region, the core promoter region (*bhsp68*) and the 3’ UTR of *Tc-hsp68* (Schinko et al., 2012).

The *pUb::lox-mYFP-lox-H2BmCherry* construct was generated by cloning the *lox-mYFP-lox-H2BmCherry* flip-out cassette under the *Tribolium pUb* element, a 940 bp DNA fragment that includes the promoter, the 5’UTR and the first intron of the *Tribolium* polyubiquitin gene (see below).

The heat-inducible Cas9 transgene (*Tc-hsp68::Cas9*) was generated by cloning the *S. pyogenes* Cas9 coding sequence under the heat-inducible promoter of the *Tc-hsp68* gene, including the endoHSE upstream region, the core promoter region (*bhsp68*) and the 3’ UTR of *Tc-hsp68* (Gilles et al., 2015; Schinko et al., 2012).

To generate the three gRNA transgenes *U6b::Lox.Uni, U6b::Lox.R* and *U6a::Lox.L*, we synthesized oligonucleotides corresponding to each gRNA targeting sequence and cloned these into p(U6b-BsaI-gRNA) or p(U6a-BsaI-gRNA), as described in Gilles et al., 2015. For *Lox.Uni* we used oligonucleotides TTCGTTCGTAATAACTTCGTATA and AAACTATACGAAGTTATTACGAA, for *Lox.R* we used oligonucleotides TTCGATCGCAATAACTTCGTATA and AAACTATACGAAGTTATTGCGAT, and for *Lox.L* we used oligonucleotides AGTGTACGTAATAACTTCGTATA and AAACTATACGAAGTTATTACGTA.

To generate the *Valcyrie.Uni* and *Valcyrie.LR* constructs, we amplified each gRNA transgene by PCR and cloned them into the plasmid carrying the *Tc-hsp68::Cas9* transgene, as illustrated in Figure 3A.

To generate transgenic lines, these constructs were cloned into the *piggyBac* transposon vector *pBac[3xP3-gTc’v]* (Addgene plasmid # 86446). The *pBac[3xP3-gTc’v]* plasmid was a gift from Gregor Bucher.

All constructs were made using a combination of conventional cloning and gene synthesis. Plasmids and annotated sequences are available at Addgene (www.addgene.org).

### Transgenic lines

Stable transgenic lines were generated as described previously (Berghammer et al., 2009) in a *Tribolium* stock of *vermilion*^*white*^ background (Lorenzen et al., 2002). Transgenic lines carrying single copies of *pBac[3xP3-gTc’v; Tc-hsp68::Cre], pBac[3xP3-gTc’v; pUb::lox-mYFP-lox-H2BmCherry], pBac[3xP3-gTc’v; Valcyrie.LR]* (transgenic line #22) and *pBac[3xP3-gTc’v; Valcyrie.Uni]* (transgenic lines #6 and #11) were established and kept as homozygous lines. The insertion of *pBac[3xP3-gTc’v; Valcyrie.LR]* in Valcyrie.LR transgenic line #39 is homozygous lethal and was therefore kept in a heterozygous state.

### *Tribolium* promoters

To identify *cis-*regulatory elements capable of driving uniform, ubiquitous expression of transgenes in early embryos of *Tribolium*, we cloned and tested the following DNA fragments: a) A 440 bp fragment of alpha tubulin, cloned by PCR using the primers TGGCCGGCCTGCAGTGAACGGTTATGATGG (including an FseI restriction site) and GGTATACTATACGAAGTTATGGTAGTTGAGTTTTACAAATTAC (including a 20 bp adaptor sequence to facilitate cloning). b) A 708 bp fragment of RPS3, cloned by PCR using the primers ATGGCCGGCCTCAACGTATGTTGTCAAACC (including an FseI restriction site) and ATGGATCCGACGTTCTAAATGGAAAAGG (including a BamHI restriction site). c) A 269 bp fragment of RPS18, cloned by PCR using the primers TTGGCCGGCCTGTCAGCGGGACATTGAC (including an FseI restriction site) and TTAGATCTGCAACCAATGCACAAACAAG (including a BglII restriction site). d) A 939 bp fragment of polyubiquitin (*pUb*), cloned by PCR using the primers TTGGCCGGCCTTTTCTTTGTCCCAAATGACC (including an FseI restriction site) and TAAGATCTGCAACGACACAAAAAATTAC (including a BamHI restriction site).

These fragments were cloned upstream of the *lox-mYFP-lox-H2BmCherry* flip-out cassette and the resulting constructs were introduced in the *pBac[3xP3-gTc’v]* transgenesis vector (see above). The activities of the *RPS3, RPS18* and *pUb* constructs were compared by observing mYFP fluorescence in *Tribolium* embryos 24 hours after injection. *RPS3* and *RPS18* gave weak fluorescence compared with *pUb*, which gave very strong fluorescence. Stable transgenic lines were then generated for the *pUb* and alpha tubulin constructs. *pUb* line 153#13 gave the strongest uniform expression and was thus chosen for subsequent experiments. We note that all the *Tribolium* ‘ubiquitous’ promoters that we have tested (including the previously published EFA fragment; Sarrazin et al., 2012) show patchy expression in late germband stages. As there are no consistent patterns, we ascribe this to noisy zygotic expression.

### Generating and screening marked clones

To generate clones using the Cre/lox approach, we crossed males homozygous for the *Tc-hsp68::Cre* transgene (transgenic line CR02) to females homozygous for *pUb::lox-mYFP-lox-H2BmCherry* (transgenic line 153#13). We collected embryos from various stages and subjected them to a heat shock for 5 min at 44°C. Embryos were then allowed to develop overnight before fixation.

To generate clones using the Valcyrie approach, we crossed males carrying the *Valcyrie.LR* or *Valcyrie.Uni* constructs to females homozygous for *pUb::lox-mYFP-lox-H2BmCherry* (transgenic line 153#13). We collected embryos from various stages (early germband to late germband elongation) and subjected them to a heat shock for 10 min at 46°C. Unless stated otherwise, embryos were allowed to develop for an additional 12 hours at 32°C before fixation. We screened the fixed embryos for native H2B-mCherry expression using a Zeiss LSM 780 confocal microscope. As a control, we screened 20 embryos from the same crosses that were collected and fixed as described above but were not subjected to a heat shock.

## Acknowledgements

We thank Manon Peleszezak and Benjamin Boumard for their work to characterize the Cre/lox system in *Tribolium*, Agnès Vallier for technical help, Peter Kitzmann and Gregor Bucher for sharing unpublished data, plasmids and transgenic lines, Vera Hunnekuhl-Terblanche, Maura Strigini, Jordi Casanova, Luke Hayden and Marco Grillo for helpful comments on the manuscript. This work was supported by a PhD fellowship from the Université Claude Bernard Lyon 1 (to A.F.G.), a post-doctoral fellowship of the Fondation pour la Recherche Médicale (to J.B.S.) and by a grant from the Agence Nationale de la Recherche [ANR-15-CE13-0014-01].

